# Composition and Diversity of CRISPR-Cas13a systems in the genus *Leptotrichia*

**DOI:** 10.1101/710533

**Authors:** Shinya Watanabe, Bintao Cui, Kotaro Kiga, Yoshifumi Aiba, Xin-Ee Tan, Yusuke Sato’o, Moriyuki Kawauchi, Tanit Boonsiri, Kanate Thitiananpakorn, Yusuke Taki, Fen-Yu Li, Aa Haeruman Azam, Yumi Nakada, Teppei Sasahara, Longzhu Cui

## Abstract

Clustered regularly interspaced short palindromic repeats (CRISPR)-Cas13a, previously known as CRISPR-C2c2, is the most recently identified RNA-guided RNA-targeting CRISPR-Cas system and has the unique characteristics of both targeted and collateral single-stranded RNA (ssRNA) cleavage activities, which was first identified in *Leptotrichia shahii* CRISPR-Cas13a. Here, the complete whole genome sequences of 11 *Leptotrichia* strains were determined and compared with 18 publicly available *Leptotrichia* genomes in regard to the composition, occurrence and diversity of the CRISPR-Cas13a and other CRISPR-Cas systems. Various types of CRISPR-Cas systems, I-B, II-C, III-A, III-D, III-like and VI-A, were unevenly distributed among the *Leptotrichia* genomes, accounting for 10 (34.4%), 1 (2.6%), 6 (15.4%), 6 (15.4%), 3 (7.7%) and 11 (37.9%) of the 29 strains, respectively, while 8 (20.5%) strains had no CRISPR-Cas system. The Cas13a effectors were highly divergent with amino acid sequence similarities ranging from 61% to 90% to that of *L. shahii* and their collateral ssRNA cleavage activities, resulting in cytotoxicity to host cell, were found to be maintained. CRISPR-Cas spacers represent a sequential achievement of former intruder encounters, and the retained spacers reflect the evolutionary phylogeny or relatedness of strains. Analysis of spacer contents and numbers among *Leptotrichia* species showed considerable diversity with only 2 (0.5%) of 400 spacers through CRIPSR-Cas I-VI were shared by two strains. The organisation and distribution of CRISPR-Cas systems (type I-VI) encoded by all registered *Leptotrichia* species revealed that effector or spacer sequences of the CRISPR-Cas systems were very divergent, and the prevalence of types I, III and VI was almost equal. There was only one strain carrying type II, while none carried type IV or V. These results provide new insights into the characteristics and divergences of CRISPR-Cas systems among *Leptotrichia* species.

## Main

### Introduction

*Leptotrichia* species, which are strict anaerobic or facultative anaerobic Gram-negative rods found in the oral cavity, intestines, female genital tract and urogenital system of both humans and animals, can ferment carbohydrates to generate multiple products, including lactic acid, formic or succinic acid, succinic acid and traces of acetic acid, depending on the substrates and species. Some *Leptotrichia* species cause opportunistic infectious diseases, including gingivitis, periodontitis, infectious endocarditis and bacteraemia (Eribe and Olsen, 2008; Eribe and Olsen, 2017). With the recent progress and spread of modern techniques, the involvement of *Leptotrichia* species in various infectious diseases has gained considerable attention (Eribe and Olsen, 2017). However, the exact clinical impact of *Leptotrichia* species relevant to infectious diseases remains unclear due to difficulties in bacterial culture, isolation and identification. To date, eight *Leptotrichia* species have been characterised, which include *L. buccalis, L. goodfellowii, L. hofstadii, L. shahii, L. trevisanii, L. massiliensis, L. hongkongensis and L. wadei.* However, many clinical *Leptotrichia* isolates have remained unclassified due to the lack of the whole-genome sequences and biochemical properties of these isolates (Eribe and Olsen, 2017). Currently, *Leptotrichia* genome data are very limited as there are only one complete whole genome sequences of species-identified strain and three unidentified strains available in the GenBank database (Gupta and Sethi, 2014; Ivanova et al, 2009).

Clustered regularly interspaced short palindromic repeats (CRISPR) and CRISPR-associated genes (*cas*) encode proteins involved in the adaptive immune response against archaea and bacteria by preventing an invasion of the host cell by foreign genetic elements, such as bacteriophages and plasmids (Marraffini, 2015). The CRISPR-Cas systems identified thus far show extreme diversity in *cas* gene compositions as well as genomic loci architecture. Despite this diversity, the latest classification system categorises the CRISPR-Cas systems into two classes based on basic compositions and structures: class 1 systems, consisting of types I and III as well as the putative type IV, which possess multi-subunit effector complexes comprised of multiple Cas proteins, and class 2 systems, which are comprised of types II, V and VI that are characterised by effector complexes consisting of a single, large Cas protein (Makarova et al., 2018; Shmakov et al., 2018). CRISPR-Cas13a (known previously as CRISPR-C2c2) is the most recently identified CRISPR-Cas system belonging to the type IV class 2 system and is characterised by cleavage activity of a single RNA-guided single-stranded RNA (ssRNA) molecule (Shmakov et al., 2015). An interesting finding recently made by Abudayyeh et al. is that CRISPR-Cas13a of *L. shahii* can perform promiscuous collateral ssRNA cleavage (Abudayyeh et al., 2016). The same research group and others recently demonstrated that the CRISPR-Cas13a system is an efficient molecular tool for the cleavage and editing of target RNA in both prokaryotic and mammalian cells (Abudayyeh et al., 2017; Abudayyeh et al., 2016; Cox et al., 2017), with highly sensitive detection of nucleic acids (East-Seletsky et al., 2017; Gootenberg et al., 2017).

The genus *Leptotrichia* is the main source of the CRISPR-Cas13a system (Shmakov et al., 2015). While the *L. shahii* CRISPR-Cas13a system has been intensively studied, the variations in the genetic features of CRISPR-Cas systems demonstrated so far have prompted us to carry out this comprehensive study of the occurrence, composition and diversity of CRISPR-Cas13a and other CRISPR-Cas systems in the genus *Leptotrichia*.

## Materials and Methods

### *Leptotrichia* strains and growth conditions

Three *L. trevisanii* strains, JMUB3870, JMUB3935 and JMUB4039, two *L. wadei* strains, JMUB3933 and JMUB3934 and one *L. hongkongensis* strain, JMUB5056, were isolated between 2012 and 2019 from blood cultures of patients in Japan. *L. wadei* strain JMUB3936 was isolated in 2017 from the bronchial lavage fluid of a patient in Japan. The following type strains of *Leptotrichia* species were obtained from the Japan Collection of Microorganisms (JCM): *L. goodfellowii* JCM16774, *L. hofstadii* JCM16775, *L. shahii* JCM16776 and *L. wadei* JCM16777. *Leptotrichia* strains were stored in Gifu Anaerobic Medium (GAM broth; Nissui Pharmaceutical Co., Ltd., Tokyo, Japan) supplemented with 40% glycerol at −80°C, and were recovered for the present study on various agar plates, of which isolates JCM12969, JCM16774, JMUB3933, JMUB3934, JMUB3935, JMUB4039 and JMUB3870 were inoculated on GAM agar, isolates JCM16775, JCM16776, JCM16777 and JMUB3936 were recovered on sheep blood agar (Kohjin Bio Co., Ltd., Saitama, Japan), and isolates JMUB5056 was grown on Anaero Columbia Agar with Rabbit Blood (Nippon Becton Dickinson Company, Ltd., Tokyo, Japan). All plates were cultured anaerobically under an atmosphere of 5% CO_2_ at 37°C for 48-72 h. *Escherichia coli* M1601 cells were grown in Luria-Bertani (LB) medium (Nippon Becton Dickinson Company, Ltd., Tokyo, Japan) with shaking or on LB agar with antibiotic supplementation (chloramphenicol at 10 μg/mL; kanamycin at 50 μg/mL) as needed for plasmid maintenance.

### Whole-genome sequencing and sequence read assembly

Bacterial genomic DNA was extracted and purified using the NucleoBond AXG kit (Takara Bio, Inc., Otsu, Shiga, Japan). DNA library preparation was performed using a Rapid Barcoding sequencing kit (SQK-RBK004; Oxford Nanopore Technologies, Oxford, United Kingdom). Sequencing was then performed using a MinION Mk-1B device integrated with a FLO-MIN-106 flow cell (Oxford Nanopore Technologies). Primary base calling was carried out using MinKNOW software (Oxford Nanopore Technologies) and the Porechop tool (version 0.2.4; https://github.com/rrwick/Porechop) was used for secondary base calling and demultiplexing of barcoded libraries. The resulting sequence reads were assembled with the Canu tool (version 1.8; http://canu.readthedocs.org/) (Koren et al., 2017). The assembled genomes were then circularised using the Circlator tool (version 1.5.5; http://sanger-pathogens.github.io/circlator/) (Hunt et al., 2015), and all of the resulting assemblies that produced contiguous sequences were polished with the nanopolish algorithm (version 0.11.0 https://github.com/jts/nanopolish) (Loman et al., 2015). Genome sequencing of all of the above strains was performed using the MiSeq platform as previously described (Watanabe et al., 2018; Watanabe et al., 2016). The draft-genomes generated by MinION were then polished again with the reads generated by MiSeq with the Pilon automated genome assembly improvement and variant detection tool (version 1.22; Broad Institute, Cambridge, MA, USA) (Walker et al., 2014). The resulting genome sequences were further trimmed using the CLC genomics workbench (Qiagen, Hilden, Germany).

### Gene Annotation and genome analysis

Gene extraction and annotation were performed with the Microbial Genome Annotation Pipeline (http://www.migap.org) or Prokka (version 1.13.3; https://github.com/tseemann/prokka) (Seemann, 2014). CRISPR-Cas genes were extracted and annotated using the CRISPRCasFinder programme (https://crisprcas.i2bc.paris-saclay.fr/) (Couvin et al., 2018). Prophages were identified by PHASTER (http://phaster.ca/) (Arndt et al., 2016). The kSNP trees were constructed with the kSNP alignment-free sequence analysis tool (version 3.021) (Gardner et al., 2015) and then visualised with the FigTree graphical viewer of phylogenetic trees (version 1.4.3; http://tree.bio.ed.ac.uk/software/figtree/). Pairwise blastn analysis for CRISPR-*cas* loci was carried out using the Easyfig application (version 2.2.2; http://mjsull.github.io/Easyfig/) (Sullivan et al., 2011).

### Plasmid construction

To examine the collateral growth inhibition activities of various Cas13a loci of *Leptotrichia* species, two sets of vectors were constructed. One set was for the expression of CRISPR-Cas13a and the other for the generation of their target RNA. The original plasmid pC003 contains a *L. shahii* Cas13a locus. First, the native Cas13a loci including native direct repeats and spacers were isolated from the genome of each *Leptotrichia* strain by polymerase chain reaction (PCR) and then cloned into the pC003 vector by replacing the *shahii*Cas13a locus using the In-Fusion HD Cloning Kit (Takara Bio, Inc., Shiga, Japan), and then kanamycin resistant gene of the vector was replaced by kanamycin-resistant gene (aphA-3) to generate pKLC-Cas13a. The pC003-LshC2C2 locus included in the plasmid pACYC184 for spacer cloning was a gift from Dr. Feng Zhang (Addgene plasmid #79152; http://n2t.net/addgene:79152;RRID:Addgene_79152).

To generate arabinose-inducible target RNA expression vector, firstly we constructed arabinose-inducible RFP expression vector (pKLC-RFP) from pC008-pBR322 by replacing tetracycline-inducible element (pBAD33) and ampicillin resistant gene (bla) to arabinose-inducible element and chloramphenicol acetyltransferase (CAT). Then the target sequence of the native Cas13a was inserted into just behind the RFP coding sequence. A native spacer sequence from each *Leptotrichia* strain and the protospacer adjacent motif, nucleotide C in this study, were adapted to the 5’ end of the reverse vector primer. The final plasmid backbone was generated by PCR followed by self-ligation using the In-Fusion HD Cloning Kit. All of the generated plasmids were confirmed by Sanger sequencing. The pC008-pBR322 with *tetR*-inducible RFP was a gift from Dr. Feng Zhang (Addgene plasmid # 79157; http://n2t.net/addgene:79157 ; RRID:Addgene_79157). The primers used in this study are listed in Supplement Table 1.

**Table 1.**
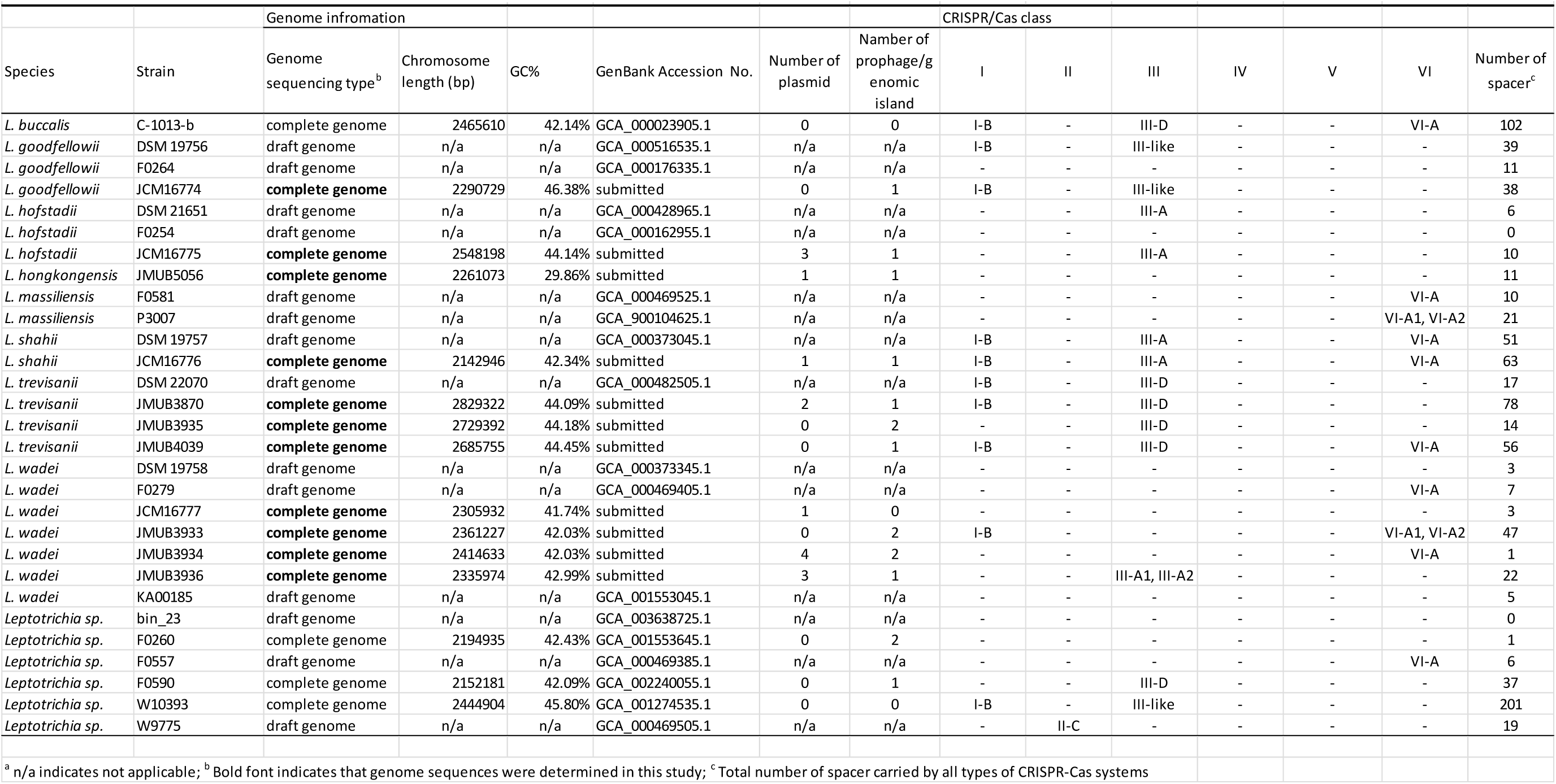
Genome and CRISPR-Cas system information of genus Leptotrichia

### Cas13a collateral growth inhibition assay

The assay for *E. coli* growth inhibition as a result of collateral ssRNA cleavage by Cas13a was carried out based on the previous paper (Abudayyeh et al., 2016). The constructed plasmids were chemically transformed into competent *E. coli* MC1601 cells. Briefly, 500 ng of CRISPR-Cas13a expression plasmid and 100 ng of arabinose-inducible target plasmid were mixed and added to the competent cells obtained above. Then, the transformation reaction was incubated on ice for 30 min, followed by heat shock at 42°C in a heat block for exactly 45 s, and then immediately transferred to tubes containing 1 mL of LB broth at room temperature, which were cooled on ice. Afterwards, the samples were incubated at 37°C for 1 h to induce antibiotic resistance and then plated on LB agar plates containing kanamycin and chloramphenicol at 37°C until colonies appeared. Then, the colonies were transferred to the LB medium containing kanamycin and chloramphenicol and incubated with gentle shaking until early stationary phase. The optical density at 600 nm (OD 600) of the bacterial culture was adjusted to 0.1 with LB medium containing kanamycin and chloramphenicol, and then 100 μl of the medium were transferred to L-shaped shaking tubes containing 10 ml of LB medium with kanamycin, chloramphenicol and 0.2 % L-arabinose to induce target gene expression. During the incubation at 25 rpm at 37°C, the optical density at 600 nm of the bacterial culture were measured every 5 minutes by TVS126MB (ADVANTEC Toyo Kaisha, Ltd.) for at least 12 hours.

### Accession of the genome sequence

The genome sequences were deposited to the DNA Data Bank of Japan (DDBJ, Bioproject accession number PRJDB7856) and the sequence raw data were deposited in the DDBJ Sequence Read Archive (DRA007809).

## Results and Discussion

CRISPR-Cas13a systems are broadly distributed among *Leptotrichia* species. The estimation of the occurrence, composition, and diversity of CRISPR-Cas13a systems in the genus *Leptotrichia* is particularly challenging because the complete whole-genome information is limited. At present, the genus *Leptotrichia* is composed of eight species. However, the complete whole genome sequences are not available except for *L. buccalis* (Ivanova et al., 2009) and three unidentified *Leptotrichia* strains. In this study, 11 strains of six *Leptotrichia* species were collected, which included *L. goodfellowii, L. hofstadii, L. hongkongensis, L. shahii, L. trevisanii* and *L. wadei*, and their complete whole-gnome sequences were determined. As it was not possible to collect isolates of the remaining species, a draft genome of *L. massiliensis* available from a public database was used. Taken together with the additional draft genome sequences retrieved from the public database, a total of 29 whole-genome sequences were used for comparative genomic analysis. All genomes were surveyed and investigated for the composition, occurrence and diversity of the CRISPR-Cas13a and other CRISPR-Cas systems. The overall features of the genomes, the distribution of the CRISPR-Cas systems, and the numbers of CRISPR-Cas spacers, plasmids and prophages for each strain are summarised in Table 1. As shown in Table 1, the chromosome size of the genus *Leptotrichia* varies from 2,142,946 to 2,829,322 bp with GC contents of 42% to 48%. Virulence factors were searched against all genomes using Prokka (version 1.13.3), but no class of genes was found to contribute to bacterial pathogenesis.

To depict the relationship among *Leptotrichia* species and strains, a whole-genome single-nucleotide polymorphism (SNP)-based phylogenetic tree was constructed of a total of 29 *Leptotrichia* genomes, which included the complete whole genome-sequencing data determined in the present study. As shown in Figure 1, four sequenced *Leptotrichia* type strains (i.e., *L. goodfellowii* DSM 19756, *L. hofstadii* DSM 21651, *L. shahii* DSM 19757 and *L. wadei* JCM16777) belonged to the same clade of the corresponding strains registered as The Department of Energy (DOE) draft-genomes (i.e., *L. goodfellowii* JCM16774, *L. hofstadii* JCM16775, *L. shahii* JCM16776 and *L. wadei* DSM 19758). The analysis also revealed that except for *L. wadei* strain JMUB3936, all strains were classified into corresponding clades. *L. wadei* strain JMUB3936 was located between *L. wadei* and *L. shahii*, and 16S rRNA sequence analysis revealed homogeneity of 97.6% with the *L. wadei* type strain JMUB3936 and 96.1% with the *L. shahii* type strain DSM19757, respectably, suggesting that strain JMUB3936 is a subspecies of *L. wadei*. Moreover, phylogenetic analysis indicated the existence of at least five unclassified *Leptotrichia* species. To date, eight species of *Leptotrichia* have been precisely described (Eribe et al., 2004; Woo et al., 2010). Apart from the eight *Leptotrichia* species, previous analysis based on the 16S rRNA gene identified 10 unclassified *Leptotrichia* species, indicating that some *Leptotrichia* strains have not yet being assigned a species name (Eribe and Olsen, 2008; Eribe and Olsen, 2017). Species classification based on both phenotype and genotype is an essential starting point to understand the biology, ecology and clinical impacts of *Leptotrichia* species.

**Figure 1.**
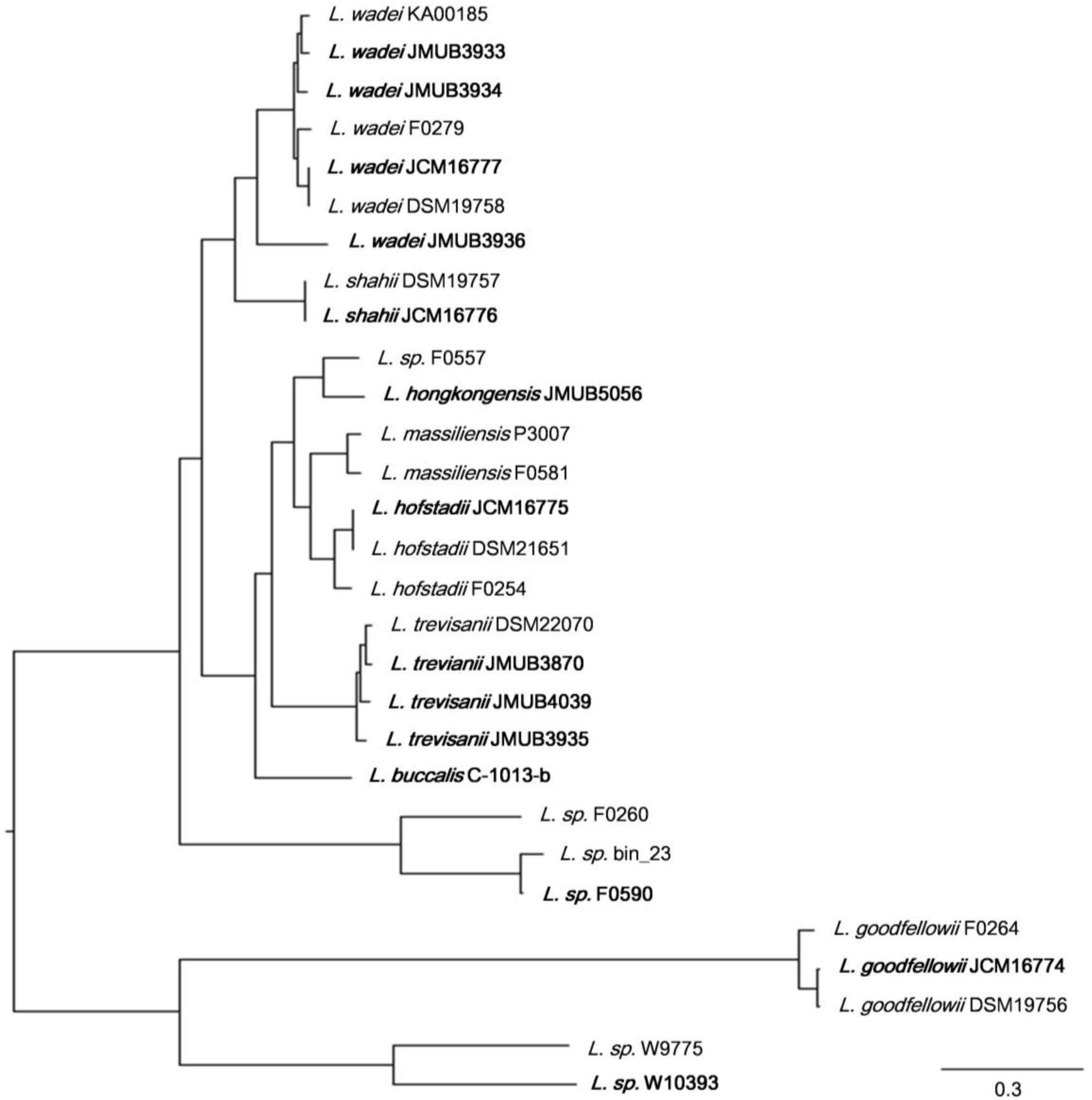
Whole genome SNP-based majority phylogenetic tree of 29 *Leptotrichia* strains generated by kSNA3.0. Horizontal branch lengths indicate changes per number of SNPs. Strains in boldface are the strains their whole genomes were determined in this study. It should be noted that this is a SNPs-based (present in at least 75% strains), not an alignment based tree and no evolutionary direction can be inferred from this tree.

To investigate the composition, occurrence and diversity of CRISPR-Cas systems among *Leptotrichia* species, all 29 *Leptotrichia* genomes were analysed for the presence of CRISPRs and *cas* genes with the use of the CRISPRCasFinder programme, which enables the identification of direct repeat consensus boundaries and the extraction of related spacers and *cas* genes (Couvin et al., 2018). This analysis identified a total of 38 CRISPR-Cas systems: 10 (26.3%) type I-B, 1 (2.6%) type II-C, 6 (15.8%) type III-A, 6 (15.8%) type III-D, 3 (7.9%) type III-like and 12 (31.6%) type VI, while no type IV and V systems were detected. Of the 29 strains, 8 (27.6%) had no CRISPR-Cas system (Table 1). Hence, we concluded that types I, III and VI were the most dominant CRISPR-Cas systems among *Leptotrichia* species.

Illustrations of all CRISPR-Cas systems identified in this study are summarised in Figure 2. Type I-B CRISPR-Cas systems were distributed in five known and one unclassified *Leptotrichia* species (Fig. 2A). It is well known that CRISPR mediates adaptive immunity according to the following three steps: (1) acquisition of short segments of foreign nucleic acids into CRISPR loci (adaptation); (2) crRNA expression and maturation (crRNA-processing); (3) target interference by recognition of the crRNA and destruction of the target nucleic acid (interference) (Bhaya et al., 2011). The *Leptotrichia* type I-B CRISPR-*cas* loci were found to include full gene sets responsible for the above three steps: *cas1, cas2* and *cas4* for adaptation, *cas6* for crRNA-processing and *cas3, cas5, cas7* and *cas8* for interference. All CRISPRs of type I-B CRISPR-Cas systems found in this study were located downstream of the *cas2* gene and/or upstream of the *cas6* gene, which was similar with a recent report of the anaerobic rod *Clostridium thermocellum* by Zöphel et al. (Ref: https://archiv.ub.uni-marburg.de/diss/z2015/0381). The type II-C CRISPR-*cas* system was found in only one *Leptotrichia* strain, W9775 (Fig. 2B). Type II-C is a class 2 CRISPR-Cas system, and the locus contains a single effector gene (*cas9*) and three adaptation genes (*cas1, cas2* and *csn2*). CRISPR was located downstream of *csn2*. Type III systems have been classified into four main subtypes (III-A to D) (Vestergaard G 2014 RNA Biol 11:156-7 and Makarova KS 2015 Nat Rev Microbiol 13:722-736). Among the *Leptotrichia* genomes analysed in this study, six type III-A, six type III-D and three type III-like loci were identified (Fig. 2C). Type III systems are reported to frequently carry accessory genes either within or immediately bordering the core *cas* gene clusters (Haft et al., 2005; Shah et al., 2019; Vestergaard et al., 2014). The type III-A systems of *Leptotrichia* can be classified into two subtypes: III-A-Lep1 and III-A-Lep2. The III-A-Lep1 loci were found in the genomes of *L. hofstadii* strain JCM16775 and *L. wadei* strain JMUB3936, which carried the full set of *cas* genes involving the adaptation, crRNA-processing and interference steps. However, the type III-A-Lep2 system found in the genomes of *L. shahii* strain DSM19757 and *L. wadei* strain JMUB3936 did not carry the *csm6* gene, which encodes an HEPN family ribonuclease that mediates sequence-unspecific ssRNA cleavage and contributes to anti-plasmid immunity (Foster et al., 2019; Niewoehner and Jinek, 2016). Instead of *csm6*, a *csx1* gene, which encodes a Csm6 homologous protein and is known to be involved in ssRNA disruption (Han et al., 2017; Sheppard et al., 2016), was identified upstream of the core *cas* genes in the type III-A-Lep2 loci.

**Figure 2.**
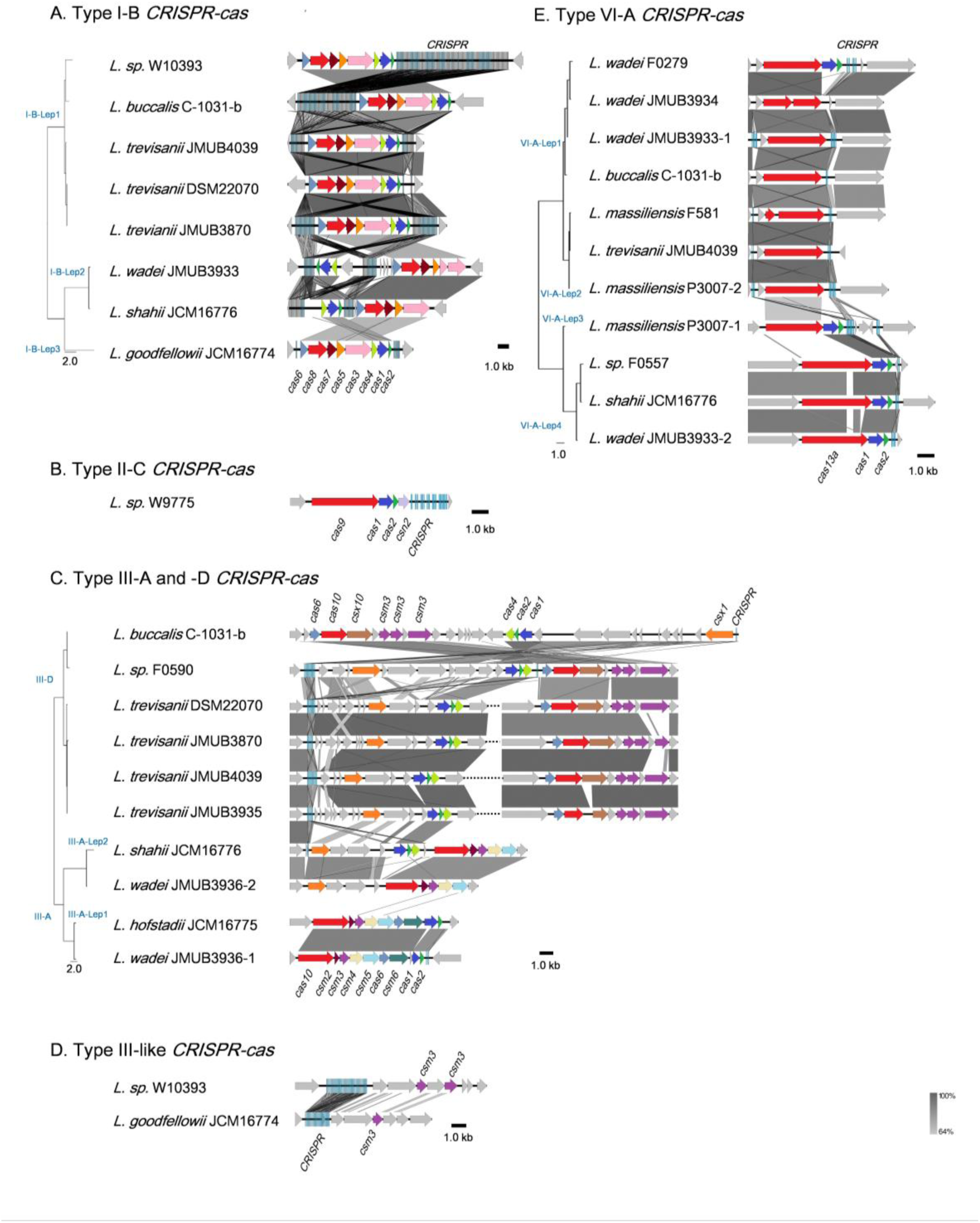
Schematic representations of the CRISPR-Cas systems in *Leptotrichia* genomes. Genetic elements of CRISPR-Cas systems identified in this study are arranged according to their relative position in the chromosome. Different color arrows indicate different categories of *cas* genes: red, genes of *cas8, cas9, cas10, cas13a*; blue, *cas1*; pink, *cas3*; yellow, *cas4*; orange, *cas5* and *csx1*; cobalt blue, *cas6*; brown, *cas7* and *csx10*; dark purple, *csm2*; purple, *csm3*; light yellow, *csm4*; light blue, *csm5*; deep green, *csm6*; gray, other genes. Left panels at the strain names represent phylogenetic parsimony trees calculated and drawn by kSNP3 and FigTree, respectively, based on DNA sequences covering all CRISPR/cas loci. Note that the regions with homologies of 64% or more show in gray bar.

*Leptotrichia* type III-D CRISPR-Cas systems were identified in four *L. trevisanii* strains, one *L. buccalis* strain and one unclassified *Leptotrichia* strain, F0590 (Fig. 2C). The genomes of *L. buccalis* strains C-1031-b and F0590, had large regions of the type III-D loci, which included core *cas* genes, *csx1*, CRISPRs and putative accessory genes. Although *L. trevisanii* strains also have type III-D gene clusters similar to those of *L. buccalis* C-1031-b and *L. sp.* F0590, the type III-D loci were split into two clusters. In *L. trevisanii* genomes, the core *cas* genes involved in crRNA-processing and interference steps were located more than 200 kb from the gene cluster of the accessory gene *csx1*, adaptation genes *cas4, cas2* and *cas1* and CRISPRs. We also found type III-like CRISPR-*cas* loci, which include *csm3* genes and CRISPRs in the genomes of *L. goodfellowii* strain JCM16774 and *L. sp.* W10393 (Fig. 2D). However, neither of these two genomes carried type III *cas* genes.

The present analysis revealed that the type VI-A CRISPR-*cas* system, which carries the single effector protein Cas13a (formerly C2c2), is broadly distributed throughout *Leptotrichia* species (Table 1). Eleven type VI-A CRISPR-*cas* loci were identified in five known and one unclassified *Leptotrichia* species. As shown in Fig. 2E, the type VI-A loci of *Leptotrichia* were further classified into four subtypes, VI-A-Lept1 to VI-A-Lept4. Cas13a is a crRNA-guided RNA-targeting effector, which has two distinct RNase motifs (O’Connell, 2019; Shmakov et al., 2015). In addition to the single RNA-guided ssRNA cleavage activity, once activated by the cognate target RNA, Cas13a becomes a promiscuous RNase that can non-sequence-specifically cleave host cell RNA (namely collateral RNA cleavage), leading to host cell death or cellular dormancy (Abudayyeh et al., 2016; East-Seletsky et al., 2016). Previously, Cas13a loci were identified and characterised in *L. wadei* F0279, *L. buccalis* C-1031-b and *L. shahii* JCM16776, which were named as LwaCas13a, LbuCas13a and LshCas13a, respectively (Abudayyeh et al.,2017; Abudayyeh et al., 2016; East-Seletsky et al., 2016). In addition, Cas13a loci were identified in three *Leptotrichia* species in the current study, *L. massiliensis, L. trevisanii* and the unclassified *Leptotrichia* strain F0557. Genomic analysis showed that, except for the cases of *L. wadei* JMUB3934 and *L. massiliensis* F581, of which *cas13a* genes were truncated, all of the VI-A CRISPR Cas13a systems identified in this study were thought to have a biological function, as based on the gene structures of the three characterised systems. Moreover, the CRISPR-associated genes *cas1* and *cas2*, encoding an adaptation module of CRISPR-Cas13a, were usually located downstream of Cas13a in subtypes VI-A-Lep3 and VI-A-Lep4. This was also the case for the VI-A-Lep1 CRISPR-*cas* system of *L. wadei* F0279. It has been suggested that the Cas13a of VI-A-Lep3 and 4, including *L. shahii* JCM16776 and *L. wadei* JMUB3933, may be more efficient than the other subtypes because the *cas13a, cas1* and *cas2* genes make up an operon with a structure that usually provides good spatial regulation during compensation.

Of the identified CRISPR-Cas13a systems, the Cas13a effectors were highly divergent with amino acid sequence similarities ranging from 61% to 90% to that of *L. shahii* JCM16776 (Table S2), which is the most intensively studied for collateral RNA cleavage activity (Abudayyeh et al., 2016). In the present study, the two most distanced CRISPR-Cas13a systems, LwaJMUB3933Cas13a-1 and LwaJMUB3933Cas13a-2, were identified in *L. wadei* strain JMUB3933. The amino acid sequence similarities of LwaJMUB3933Cas13a-1 and LwaJMUB3933Cas13a-2 to that of *L. shahii* JCM16776 (LshCas13a) were 61% and 87%, respectively. In addition, Cas13a was firstly identified in *L. trevisanii* genome (*L. trevisanii* JMUB4039, LtrCas13a). The cytotoxic activities dependent on collateral ssRNA cleavage activity of the four systems (LwaJMUB3933Cas13a-1 and -2, LtrCas13a, LshCas13a) were compared with the use of a co-expression assay, where two different vectors carrying the CRISPR-Cas13a with spacer sequence and target sequences were co-transformed into competent *E. coli* cells, and then the growth curves were observed with or without 0.2% arabinose inducing mRNA which contain the target sequences. As shown in Figure 3, except for spacer-1 targeting of LwaJMUB3933Cas13a-1, the four Cas13a systems could significantly affect the growth of the host cells, indicating that the collateral cellular dormancy as a result of ssRNA cleavage were maintained among those strains. Here we demonstrated the collateral cytotoxic activities of four Cas13a using representative native spacers, a comprehensive study is necessary to draw precise conclusions regarding enzymatic activities.

**Figure 3.**
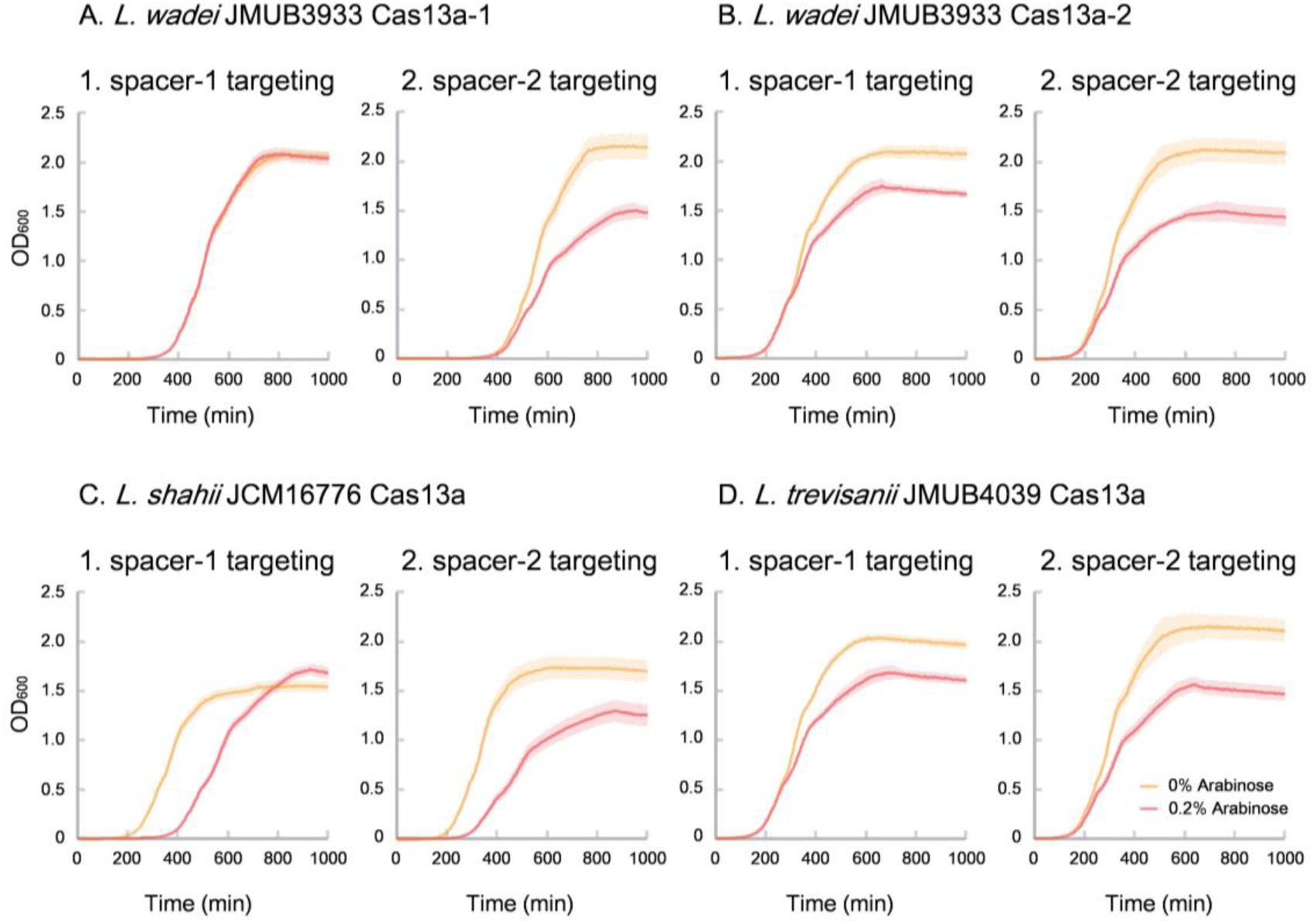
Collateral growth inhibitory activities of LwaCas13a-1, LwaCas13a-2, LtrCas13a and LshCas13a. Native CRISPR-Cas13a expression vector carrying kanamycin-resistant gene and arabinose-inducible target RNA (protospacer) expression vector which carries chloramphenicol-resistant gene were co-transformed into E. coli MC1061 and incubated at 37°C on LB plates containing chloramphenicol and kanamycin until colonies appeared. The independent colonies were transferred to LB medium containing chloramphenicol, kanamycin and 0.2% arabinose (inducer of target RNAs) and incubated with gentle shaking at 25 rpm at 37°C. Optical density at 600 nm (OD 600) were measured every 5 minutes and the growth curves were obtained.

CRISPR-Cas spacers represent a sequential achievement of former intruder encounters, and the retained spacers reflect evolutionary phylogeny or relatedness of the strains. In the present study, the CRISPR spacers of all strains were analysed, as well as the protospacers of the strains with complete whole genomes. A protospacer is a spacer homologue existing in the plasmid, phage or chromosomal DNA that matches the spacers of CRISPR elements. The number and distribution of spacers and the corresponding protospacers are summarized in Table S3 and Figure 4. As shown in Figure 4, the number of spacers had varied among the strains, ranging from 1 to 157, which reflects the biological diversity of *Leptotrichia* species that acquired a CRISPR-Cas system for mediation of immunity. The protospacer distribution showed that cross-targeting events within the species were very rare, but often occurred inter-species. Notably, some spacers matched against protospacers existed in the same genome. As a possible interpretation, CRISPR-Cas systems may be regulated by unknown mechanisms that degrade their own function, as self-targeting results in cell death.

**Figure 4.**
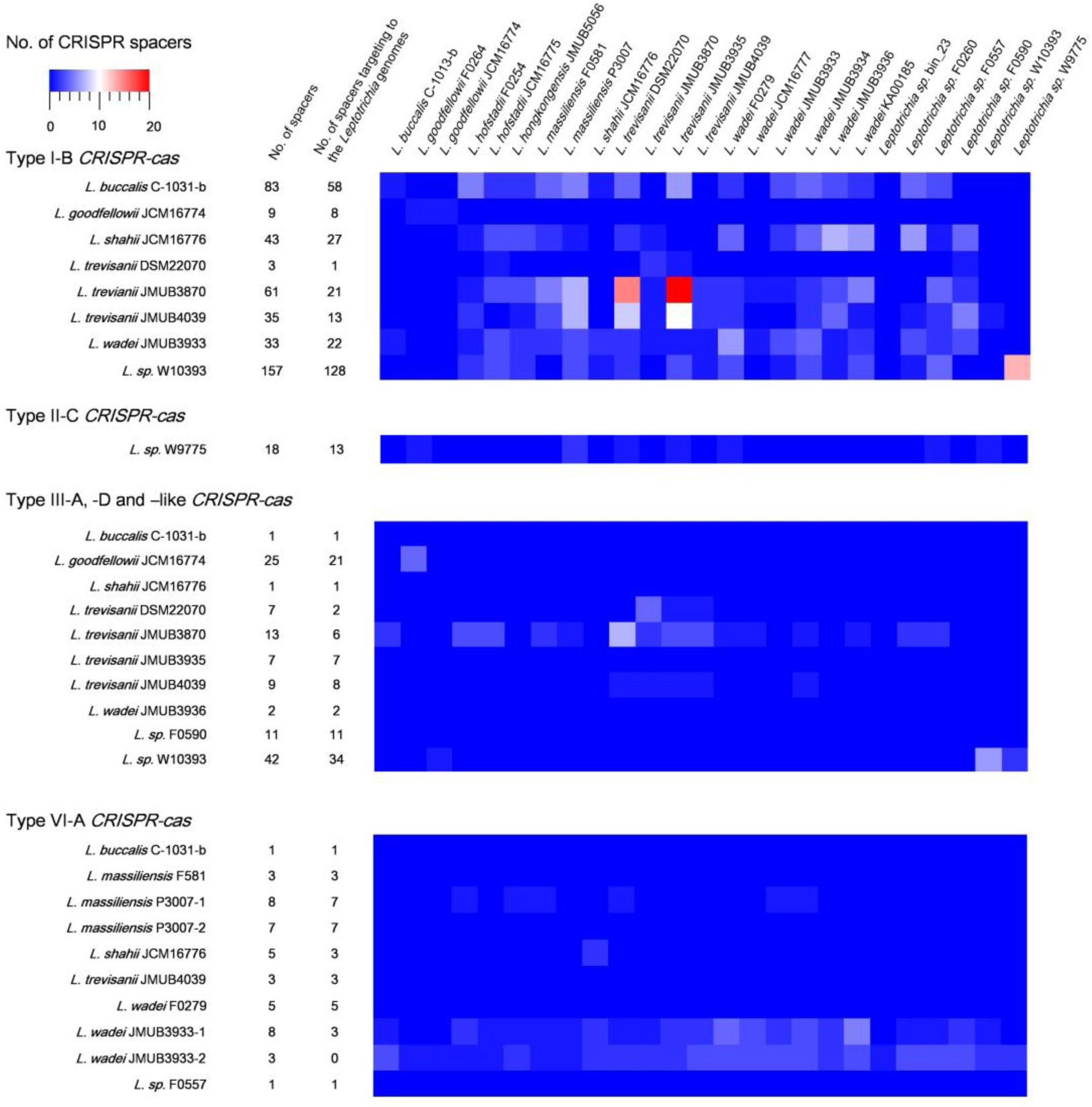
Match of *Leptotrichia* CRISPR-Cas spacer sequences with sequences of phages, plasmids and bacterial genomes of own bacterial genus of *Leptotrichia*. This hit map shows number of spacers and the present of matched sequence with homology at least 90% in genomes of genus *Leptotrichia*. The first left panel indicates a total numbers of spacers present in each strain, and the second left panel does the numbers of spacer which have hit matches against each strain.

In summary, the complete whole-genome sequences of 12 *Leptotrichia* strains and the CRISPR-Cas systems of eight *Leptotrichia* species were determined. The prevalence of the CRISPRE-Cas systems was determined in all 29 genomes available in public databases. The results showed that 21 (72.4%) strains carried CRISPRE-Cas systems, and types I, III and VI were predominantly distributed among *Leptotrichia* species, but not types IV and V. The presence of a CRISPR-Cas system in more than 72% of the genomes investigated suggests a meaningful role of this system in the biology of *Leptotrichia* species.

## Data Availability

The genome sequence has been deposited at DDBJ/GenBank: PRJDB7856 (Bioproject) and DRA007809 (Raw data).

## Conflict of Interest Statement

The authors declare that the research was conducted in the absence of any commercial or financial relationships that could be construed as a potential conflict of interest.

## Acknowledgement

We thank Dr. Toyoko Oguri at Juntendo University, Dr. Katsuko Okuzumi at Dokkyo University Hospital, Dr. Aki Kawashima, Dr. Haruki Sawamura, Dr. Akitaka Yokoyama at Ichinomiyanishi Hospital, Dr Noriyuki Abe at Tenri Hospital, and Dr. Eri Morita, Dr. Yuji Hatano and Dr. Satoru Hayashi at Japanese Red Cross Gifu Hospital for kindly providing *Leptotrichia* strains. We also thank Dr. Feng Zhang at Massachusetts Institute of Technology and Dr. Masato Suzuki for kindly gifting plasmids.

## Funding

This work was supported by JSPS KAKENHI (Grant No. 15H05654 and 19K08960 to SW, 18K15149 to KK, 17K15691 to YS, 19K15740 to MK and 17K19570 to LC), the Takeda Science Foundation (SW, LC), the JSPS International Research Fellow (Grant No. 17F17713 to LC) and the Japan Agency for Medical Research and Development J-PRIDE (grant no. JP19fm0208028 to LC). The funders had no role in the study design, data collection and analysis, decision to publish or preparation of the manuscript.

## Authors’ contributions

SW, KK and LC designed the study. SW, BC, KK, YA, XET, YS, MK, TB, KT, YT, FYL, AHA and TS performed experiments and analyzed data. YN coordinate the clinical sample collection. SW, BC and KK prepared figures and drafted the manuscript. SW, BC, KK, XET and LC edited and revised the manuscript. All authors have read and approved the final manuscript.

---

**Table 1** *Leptotrichia* strains used in this study

**Table S1** Primer list

**Table S2** amino acid sequence identities of Cas13a in Leptotrichia

**Table S3** CRISPR and spacer sequences in Leptotrichia genomes

